# *Bacillus toyonensis* biovar Thuringiensis: an overlooked entomopathogen?

**DOI:** 10.1101/2024.07.21.604280

**Authors:** Diego Herman Sauka, Cecilia Peralta, Eleodoro E. Del Valle, Leopoldo Palma

## Abstract

Horizontal gene transfer (HGT) significantly influences prokaryotic genome evolution. *Bacillus cereus* and *Bacillus thuringiensis* are nearly identical at the chromosomal level, except for *B. thuringiensis* producing parasporal crystals. The genes for these crystal proteins (e.g., *cry1A*), along with other encoded insecticidal proteins (e.g., *vip3A*), are located on megaplasmids and can be horizontally transferred. Recently, Sauka et al. (2022) reported a *Bacillus toyonensis* strain that produces parasporal crystals with dual insecticidal activity. This strain was classified as *Bacillus toyonensis* biovar Thuringiensis (NCBI: txid2923195) following Carroll et al.’s (2020) nomenclature. Misclassified *B. toyonensis* strains, previously identified as *B. thuringiensis* (e.g., strain MC28), encode *cry* and *cyt* genes toxic to lepidopteran and dipteran insects. Advances in genome sequencing and bioinformatics tools now reduce misidentifications, enabling accurate reclassification in databases like GenBank. These findings highlight the need for genome-based taxonomic reassessment within the *Bacillus cereus* group and clarify the chromosomal placement of crystal-forming *B. toyonensis* strains.

---

*Bacillus thuringiensis* is a gram-positive, spore-forming bacterium known for producing various proteins with specific biocidal activity against invertebrates. This characteristic has made *B. thuringiensis* one of the most studied and widely used biological control agents for sustainable pest management in agriculture (targeting insects and nematodes) and mosquito control (1). The repertoire of pesticidal proteins includes parasporal crystal proteins such as Cry, Tpp, and Cyt (δ-endotoxins), produced during the stationary growth phase, and secreted proteins historically referred to as vegetative insecticidal proteins (Vip), including the binary Vpb1/Vpa2 (formerly Vip1/Vip2), Vip3, and Vpb4 (formerly Vip4) (2, 3). Vip3 proteins can be produced from mid-log growth phase through sporulation, with *vip3A* transcription activated at the onset of stationary phase by the regulator VipR (4).

Other gram-positive, spore-forming bacteria also produce biocidal proteins. For example, *Lysinibacillus sphaericus* produces Tpp proteins (formerly BinA/BinB proteins) (5), and *Brevibacillus laterosporus* produces Cry proteins, both of which exhibit toxicity against mosquitoes (6).

In 2022, Sauka et al. isolated and preliminarily classified a *B. thuringiensis* strain with parasporal crystals and insecticidal activity against *Cydia pomonella* (Lepidoptera) and *Anthonomus grandis* (Coleoptera) (7). However, with an average nucleotide identity (ANI) of 98.6% against *Bacillus toyonensis*, this novel strain was officially named *B. toyonensis* biovar Thuringiensis strain Bto_UNVM-94 (7), following the nomenclature system proposed by Carroll et al. (2020) (8). Another similar case is reported for *Bacillus wiedmannii* biovar thuringiensis, a strain harboring genes encoding for parasporal crystal proteins with mosquitocidal activity (9). *B. toyonensis* was first described in 2013 by Jimenez et al. as a member of the *Bacillus cereus* group, which includes *B. cereus, B. thuringiensis*, and other related pathogens. This species has been used as a probiotic in the swine, poultry, cattle, rabbit, and aquaculture industries for over 30 years (10).

Samples were taken with a tubular soil sampler from different areas of Argentina, namely: a native forest area from Jujuy province (strain Bto_UNVM-42), Tancacha town from province of Córdoba (strain Bto_UNVM-100) and as previously described for strain Bto_UNVM-94 (7). After collection, samples were placed in zip-lock bags and stored at 4 °C until bacterial isolation could be performed. For this purpose, 3 grams of the soil were suspended in 10 ml of sterile distilled water and homogenized by vortexing for 1 minute. The suspension was then subjected to a heat shock at 80 °C for 30 minutes to select for spore-forming bacteria. Serial decimal dilutions (from 10^−3^ to 10^−5^) were prepared, and 50 μl aliquots from each were plated on nutrient agar medium composed of 0.5% peptone, 0.3% beef extract, 0.5% NaCl, and 1.5% agar. The plates were spread using a Drigalsky spatula and incubated at 28 °C for 48 to 72 hours. Colonies resembling *Bacillus thuringiensis*, characterized by a matte-white appearance and irregular margins, were selected and further streaked for purification on fresh agar plates. These were incubated under the same conditions until sporulation was observed. Once sporulated, cultures were heat-fixed on glass microscope slides and stained with a Coomassie Blue solution (0.133% in 50% acetic acid) for visualization in a light microscope (11). The presence of parasporal crystals, characteristic of a *B. thuringiensis*-like phenotype, was later confirmed using scanning electron microscopy.

Genomic DNA, encompassing both chromosomal and plasmid components, was extracted using the Wizard Genomic DNA Purification Kit (Promega). The procedure was carried out in accordance with the manufacturer’s protocol, specifically optimized for DNA isolation from gram-positive bacterial strains. The extracted DNA was used subsequently for the preparation of a pooled Illumina library at the Genomics Unit from the National Institute of Agricultural Technology (INTA, Argentina) by using high-throughput Illumina sequencing technology for strains Bto_UNVM-42 along with Bto_UNVM-100, and as previously described for strain Bto_UNVM-94 (7). The raw Illumina reads obtained were first trimmed and assembled into contigs by using Velvet plug in included in Geneious R11 (www.geneious.com). The resultant contigs were then analyzed with BLAST (12) using a customized non-redundant insecticidal protein database. Genome annotation was performed with the NCBI Prokaryotic Genome Annotation Pipeline (2021 release) although it was also annotated with RAST (13). Species delimitation was performed with the Type (Strain) Genome Server (14), FastANI (15) and with PubMLST (16).

By using this combination of genome-genome hybridization analysis with TYGS server (14) and % ANI calculations with FastANI v0.1.3 (15), we have detected the potential misclassification of two *Bacillus* species stored in the GenBank database (17). Strains *B. thuringiensis* MC28 (RefSeq GCF_000300475.1) (18) and *B. cereus* Rock 1-3 (RefSeq GCF 000161155.1) (19) are clustered in the same branch along with type strain *B. toyonensis* P18 (RefSeq GCF_016605985.1) (Figure 1).

**Figure 1.**
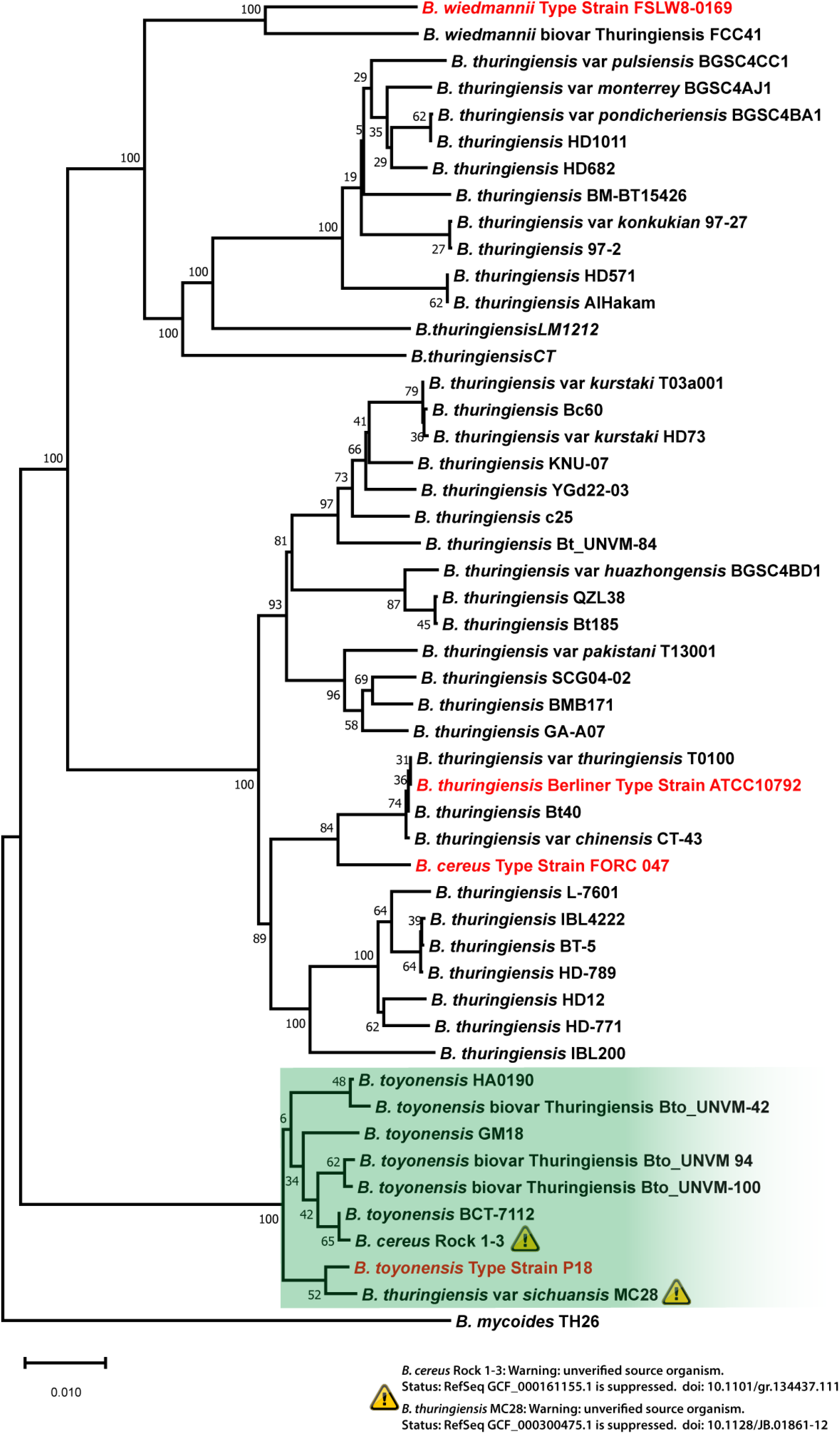
Genome BLAST Distance Phylogeny (GBDP) [18] based on genome data (whole-genome sequence-based) using TYGS server. The *B. toyonensis* cluster is highlighted in green color. The exclamation marks indicate potentially misclassified *B. toyonensis* strains.

In addition, estimated % ANI and PubMLST support values of these strains, strongly suggests the need of performing a reclassification of these *Bacillus* species by following the novel nomenclature system proposed by Carroll et al., (2020). We also added in the calculations the genomic sequences coming from novel *Bacillus* strains Bto_UNVM-42 and Bto_UNVM-100 (Table 1).

**Table 1.**
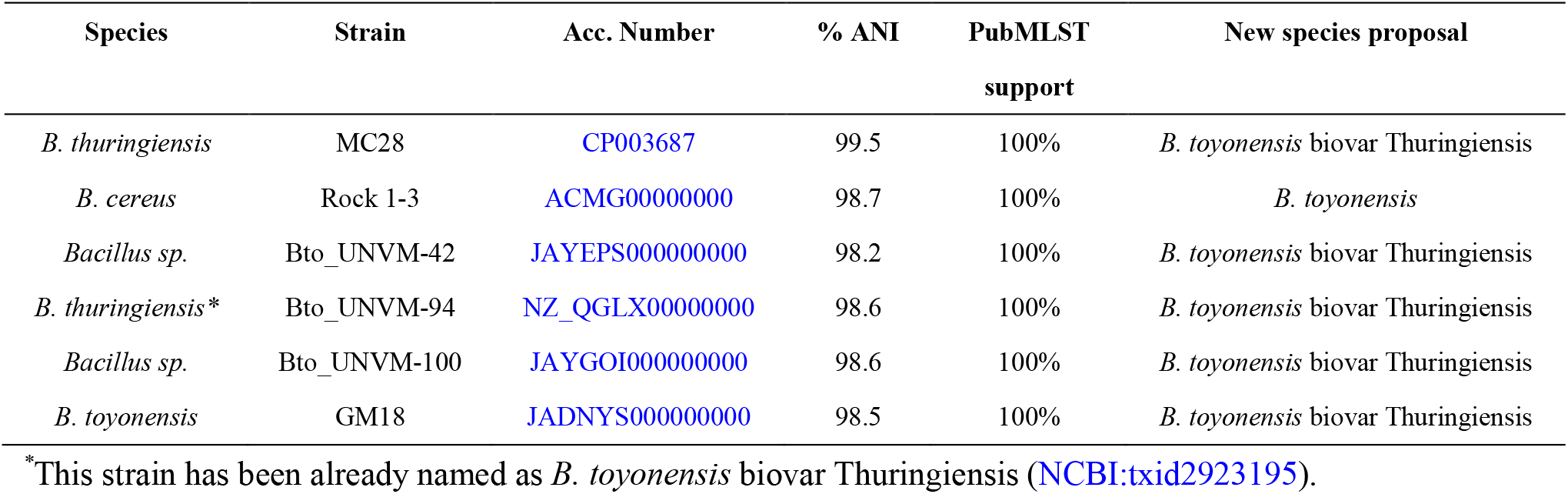
% ANI calculation estimated by comparison with type strain *B. toyonensis* P18, showing the reclassification proposal following the nomenclature system by Carroll et al., (2020). A 95-96% ANI is generally accepted as the species boundary (28), therefore, genomes that share 95-96% ANI or higher may be considered as belonging to the same species.

Strain MC28 can form *B. thuringiensis*-like parasporal crystals and harbor several genes encoding for δ-endotoxins (Cry and Cyt), reported to be highly toxic to lepidopteran and dipteran insects, according to the original strain description (18). However, this strain, which clusters with *B. toyonensis* in genome-based phylogenetic analyses, has gone unnoticed due to its original classification as *B. thuringiensis* MC28. We hypothesize here the existence of horizontal gene transfer (HGT) events. HGT is one of the most significant sources of evolution that continuously causes major changes into prokaryotic genomes and has been previously described within the *Bacillus* genus (20). One consistent example of HGT within the *Bacillus cereus* group is the transfer of genes encoding insecticidal proteins, such as the parasporal crystal proteins (Cry and Cyt), which confer toxicity against insect pests. These genes are predominantly plasmid-borne and are frequently found in genomic regions associated with insertion sequences and other mobile elements, which contribute to their horizontal dissemination (3, 20-23). Several studies have documented the occurrence of HGT events involving insecticidal genes within the *B. cereus* group. For instance, Liu et al. (2015) investigated the genomic diversity of *B. thuringiensis* strains and identified multiple instances of HGT-mediated acquisition of insecticidal genes from other *Bacillu*s species (24). Similarly, Geng et al. (2023) demonstrated HGT of the 144-kb plasmid pTAND672-2, carrying mosquitocidal toxin genes, through conjugation experiments, between *B. thuringiensis* serovar *israelensis* and *L. sphaericus* (25).

Overall, HGT serves as a key mechanism driving genetic diversity and adaptation within the *B. cereus* group, particularly in the context of insecticidal properties.

In addition, the *B. cereus* strain Rock 1-3 has been also potentially misclassified and should be renamed as belonging to *B. toyonensis* species.

In the case of *Bacillus* sp. strains Bto_UNVM-42 and Bto_UNVM-100, the same hypothesis involving HGT events may apply. Strain Bto_UNVM-42 and Bto_UNVM-100 have been isolated from provinces of Jujuy and Córdoba (Argentina), respectively, and they were able to produce parasporal crystals with their genomic sequences harboring several genes with potential insecticidal activity (Table 2). However, both TYGS server and estimation of % ANI against type strain *B. toyonensis* P18 showed these strains as belonging to *B. toyonensis* species. Following the nomenclature proposed by Carroll et al., (2020), we propose to name these strains as belonging to *B. toyonensis* biovar Thuringiensis species.

**Table 2.** Putative insecticidal gene content of *B. toyonensis* strains.

| Strain | Homolog Proteins | Potential targets | Parasporal crystals | References |
| --- | --- | --- | --- | --- |
| MC28 | Cry4C, Cry30F, Cry53A, Cry54A, Cry68A, Cry69A, Cry70B, Cyt1D, Cyt2A | Lepidoptera and Diptera | Spherical | (18) |
| Bto_UNVM-42 | Cry32E, Cry32G, Cry32K, Cry32M, Cry32U, Cry69A, Cry73B, Cyt1A, Cyt1D, Mpp3A, Mpp23A | Lepidoptera, Coleoptera and Diptera | Bipyramidal | This work |
| Bto_UNVM-94 | Cry7G, Mpp2A | Lepidoptera and Coleoptera | Bipyramidal | (7) |
| Bto_UNVM-100 | Cry7B, Cry7G, Cry73A, Mpp2A, Xpp22A | Lepidoptera and Coleoptera | Bipyramidal | This work |
| GM18 | Xpp22A, App4A, Tpp78A, Vpb1A, Mpp75A, Vpb1E | Human-cancer cells | Produced, morphology is not specified | (26) |

Following the same approach, our results indicate that strain GM18 has been correctly classified as *B. toyonensis* (26). Based on the reported production of parasporal inclusions and the presence of pesticidal and parasporin-related genes in its genome, we propose that this strain be designated *B. toyonensis* biovar Thuringiensis, consistent with the nomenclature framework proposed by Carroll et al. (2020). Although GM18 was reported to lack insecticidal activity, solubilized parasporal inclusion proteins were shown to display cytotoxic activity against human cancer cell lines in the original description (24) (Table 2). Based on our phylogenetic analyses, the *B. toyonensis* biovar Thuringiensis strains described here exhibit phenotypic characteristics like those of *B. thuringiensis*. These strains not only produce parasporal crystals but also exhibit biocidal activity against insects and human cancer cells.

In this work, we provide genomic and taxonomic evidence indicating that several *Bacillus* strains previously classified as *B. thuringiensis* or *B. cereus* in fact belong to *B. toyonensis*, and that a subset of strains of the latter species harbor genes encoding parasporal crystal proteins (Cry and Cyt) and other pesticidal factors (Vpb, App, Mpp and Tpp proteins) (27). These findings support the recognition of crystal-forming *B. toyonensis* strains as part of the taxonomic and phenotypic diversity within the *B. cereus* group. Our hypothesis that HGT events have contributed to the acquisition of *cry, cyt* and other insecticidal genes provides a plausible explanation for the emergence of a *B. thuringiensis*-like phenotype in strains that are genomically assigned to a different species. This observation is consistent with the well-documented role of mobile genetic elements in shaping ecological traits within the *B. cereus* group. Taken together, these observations highlight the importance of genome-based taxonomic frameworks for resolving species boundaries within the *B. cereus* group. The identification of crystal-forming strains that are chromosomally assigned to *B. toyonensis* illustrates how accessory gene content may complicate phenotype-based classification and supports the use of genome-informed nomenclature.

Future studies should aim to clarify the frequency, directionality and ecological drivers of these HGT events in natural soil populations, as well as to determine how widespread *B. thuringiensis*-like phenotypes are within *B. toyonensis* and related taxa. Such efforts will be essential to fully appreciate the evolutionary dynamics, ecological significance and biotechnological potential of mobile pesticidal gene reservoirs within the *B. cereus* group.

## Acknowledgments

Leopoldo Palma would like to express gratitude to the Spanish Government-Universities Ministry, the Next Generation EU and Recovery, Transformation, and Resilience plans for funding his awarded María Zambrano contract (ref. ZA21-003) and the Ramón y Cajal programme (Ref. RYC2023-043507-I).

